# Glutathione Peroxidase 8 (GPX8)-IL6 axis is essential in maintaining breast cancer mesenchymal stem-like state and aggressive phenotype

**DOI:** 10.1101/818245

**Authors:** Anees Khatib, Solaimuthu Balakrishnan, Michal Ben-Yosef, Gidi Oren, Areej Abu Rmaileh, Michal Schlesinger, Jonathan H. Axelrod, Michal Lichtenstein, Yoav D. Shaul

## Abstract

Metabolic reprogramming as a downstream result of oncogenic signaling pathways has been described as a hallmark of cancer. Here, we describe a reverse scenario in which a metabolic enzyme regulates cancer cell behavior by triggering a signaling pathway. We find that glutathione peroxidase 8 (GPX8), a poorly characterized redox enzyme that resides in the endoplasmic reticulum, is upregulated during the epithelial-to-mesenchymal (EMT) program in HMLE and A549 cells. In cancer patients, high tumor levels of GPX8 correlate with mesenchymal markers and poor patient outcome. Strikingly, GPX8 knockout in mesenchymal-like cells results in an epithelial-like morphology, downregulation of EMT characteristics, loss of cancer stemness features, and impeded tumor initiation in mice. We determine the mechanism governing this reduction in cancer aggressiveness is through the repression of crucial autocrine factors, in particular, interleukin-6 (IL-6). Specifically, GPX8 knockout impairs IL-6-driven activation of the JAK-STAT3 signaling pathway, a critical regulator of a cancer-aggressive state. Altogether, we uncover the GPX8-IL-6 axis as a novel metabolic-inflammatory pathway that acts as a robust EMT activator and program to induce aggressive cancer cell characteristics.

## Introduction

Over the past few decades, intensive studies in breast cancer have led to the development of diagnostic technologies for early disease detection and selective treatment for specific tumor subtypes. However, despite these advances in breast cancer therapy, many patients still develop tumor metastases due to disease recurrence and therapeutic resistance (1). One of the proposed mechanisms by which breast carcinoma cells escape the effects of cancer treatment is cancer cell trans-differentiation into mesenchymal-like cells, which is mediated by a cellular program called epithelial-mesenchymal transition (EMT) (2). This cellular transition is accompanied by significant changes in cell morphology, resulting in the elimination of cell-cell interaction and the loss of cell polarity. These alternations explain how the carcinomas gain the ability to detach from the primary tumor and promote the metastatic cascade (3).

The EMT program is regulated through the activation of a core set of EMT-activating transcription factors (EMT-TF) (4). The activity of these EMT-TF regulated by intracellular signaling pathways that are induced by specific ligands such as chronic transforming growth factor-beta (TGFβ), WNT signaling pathway, NOTCH pathway, mitogenic growth factors (1), and inflammatory cytokines, such as IL-6 (5, 6). In recent years many studies demonstrated a link between the EMT program and cancer stemness cells (CSC) (7, 8). Induction of the EMT program in epithelial cells results in the expression of stemness markers such as CD44^high^/CD24^low^ and the ability to form mammospheres (8–10). However, these CSC are not the outcome of the full execution of the EMT program, but instead are at an intermediate state along the epithelial-mesenchymal spectrum (1).

Tumors exhibit a unique cellular metabolism pattern relative to non-proliferative normal cells, in that they need to satisfy the enhanced cell growth and proliferation (reviewed in (11–13)). However, cancer-dependent metabolic rewiring was found to be more complicated than initially described (14), since in addition to a generic metabolite demand, it became evident that there are also metabolic enzymes that are essential only for a certain types of tumors (15–18). These findings imply that the demand for metabolic rewiring in promoting malignancy is not limited only to support cell replication but is also required in order to satisfy other cellular needs. Thus, the cancer-dependent metabolic rewiring plays an essential role in promoting proliferation-independent processes such as supporting and maintaining the gain of traits associated with high-grade malignancy. To systematically identify the metabolic enzymes which regulate tumor progression, we generated MERAV, a web-based tool to analyze human gene expression between different cancer types and normal tissues ((http://merav.wi.mit.edu/,(19)).

Recently we characterized a set of 44 metabolic genes that are selectively present in high-grade tumors bearing mesenchymal markers, that we designated as “mesenchymal metabolic signature” (MMS) (10). The pattern of expression of the MMS genes suggests their critical role in acquiring the high-grade malignancy features. In order to systematically determine the role of MMS, we developed a FACS-based shRNA screen that identified 16 genes as essential for the EMT program (10). Among them is dihydropyrimidine dehydrogenase (DPYD), the rate-limiting enzyme of the pyrimidine degradation pathway (20), whose activity, as has been demonstrated both *in vivo* and *in vitro*, is essential for carcinoma cells to gain mesenchymal characteristics (10). The finding of DPYD essentiality for the EMT program motivated us to determine the role of the glutathione peroxidase 8 (GPX8) another screen-hit MMS gene.

The primary function of the glutathione peroxidase (GPx) family of proteins is to limit the cellular accumulation of the reactive oxygen species (ROS) (21). These enzymes use glutathione (GSH) as reductant (22), catabolized peroxides to the corresponding alcohols. The unique activity of GPx is determined by their particular amino acid composition, as most members are selenoproteins with selenocysteine in the catalytic center (GPX1-4, and GPX6). (22). GPX8, the last member of this family to be identified (23), is a type II transmembrane protein with high sequence similarity to the soluble GPX7 (NPGPx). Both enzymes share many characteristics as they contain a KDEL-like endoplasmic reticulum (ER) retrieval motif and are therefore localized in ER (24) and in ER- mitochondria-associated membranes (MAM) (25). Despite their name and their similarity to the other members of the GPx family, both GPX7 and GPX8 have low GPx activity (24), as they lack the GSH-binding domain (26). The proposed function of both GPX7 and GPX8, in association with protein disulfide isomerase (PDI) peroxide-mediated oxidative protein folding (24), is to specifically bind and clear the peroxides generated in the ER by the endoplasmic reticulum oxidoreductase 1 Alpha (ERO1α) enzyme, which introduces disulfide bonds into PDI (27). The physiological function of GPX8 is still unclear, as it has been reported to be involved in diverse physiological processes. For example, its serves as a cellular substrate to the hepatitis C virus NS3-4A protease (28), and ectopic expression of GPX8 in the rat pancreatic β-cells induces ER stress (29). Our knowledge about GPX8 regulation is still limited, as its expression was found to be regulated by hypoxia-inducible factor (HIF1α) (30) and repressed by insulin-like growth factor 1 receptor (IGF1R) in the lung (31). These studies are only starting to reveal some physiological roles of GPX8, but its function in the biology of tumors is still unclear. Here we report that GPX8 robustly regulates the EMT program via the regulation of IL-6-trans-signaling, mediated by the soluble IL6 receptor that results in the inhibition of the JAK-STAT3 signaling pathway.

## Results

### GPX8 expression correlates with mesenchymal markers

Global gene expression analysis identified the MMS gene GPX8 to be upregulated in cells bearing mesenchymal markers (10). We therefore first validated GPX8 expression in large-set gene expression databases of cancer cell lines. Previous unsupervised hierarchical clustering analysis of MERAV (http://merav.wi.mit.edu/)(19) determined that the ~1,000 different cancer cell lines segregated into five distinct groups based on their metabolic gene expression (10). These groups include a set of 378 cell lines originating from epithelial tissues (epithelial group) and a set of 150 cell lines that share the expression of mesenchymal markers (mesenchymal group). We found GPX8 expression to be significantly upregulated in mesenchymal relative to epithelial cell lines. This expression pattern is similar to the other known mesenchymal markers such as, DPYD (10), fibronectin (FN1) (9), ZEB1 (32), ZEB2 (33) and cadherin 11 (CDH11) (34) (Fig. 1A). In contrast, the expression of the epithelial markers cadherin 1 (CDH1 (E-cadherin)) (32), claudin (CLDN1) (9) and junction plakoglobin (JUP, (γ-catenin) anticorrelates with GPX8 expression pattern.

**Figure 1:**
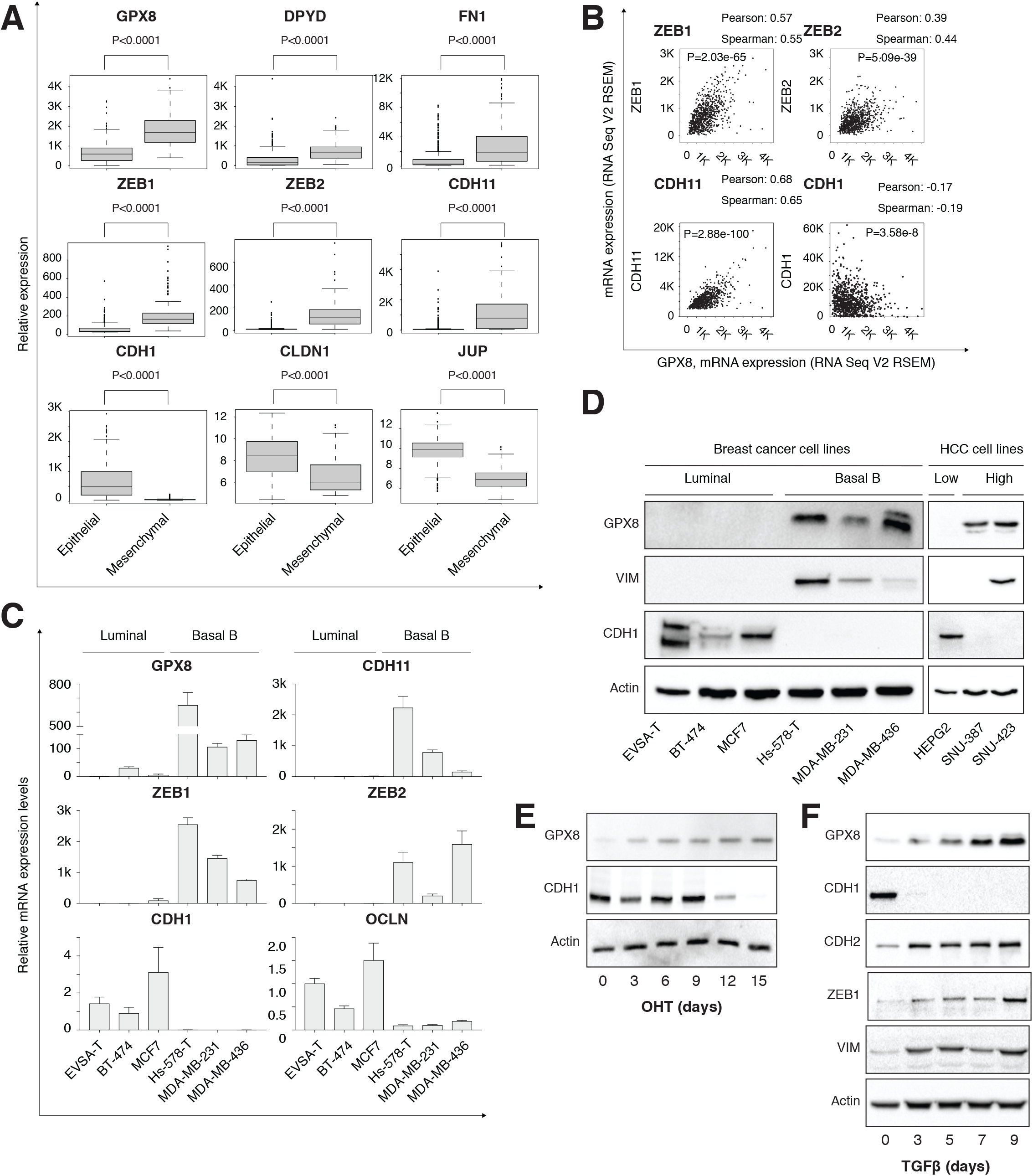
GPX8 expression is upregulated in mesenchymal-like cells: (**A**) Elevated GPX8 gene expression in mesenchymal cell lines. Cancer cell lines were divided into epithelial (378 cell lines) and mesenchymal (150 cell lines) groups based on the expression of known mesenchymal markers. Box plots represent the expression levels of the indicated genes in each group. P value was determined by Student’s T-test using R. (**B**) GPX8 expression correlates with mesenchymal markers. Patients gene expression data was generated by the TCGA project and analyzed using the cBioportal webtool. GPX8 expression positively and significantly correlates with known mesenchymal markers (ZEB1, ZEB2, and CDH11) and negatively with the epithelial marker CDH1. The Pearson, Spearman correlation and the P value were calculated using the cBioportal website (https://www.cbioportal.org). (**C**) GPX8 mRNA level is upregulated in basal B breast cancer cells. The relative level of GPX8, as well as other indicated EMT markers in breast cancer cell lines, were determined by quantitative real-time PCR (qPCR). The expression level of all cell lines is relative to that of the EVSA-T cell line. Each value represents the mean ± SD for N=3. (**D**) Individual validation of GPX8 protein levels in the indicated breast cancer and HCC cell lines. Cells were lysed and subjected to immunoblotting using the indicated antibodies (**E**) GPX8 expression is upregulated in HMLE-Twist-ER inducible system. HMLE-Twist-ER cells were treated with hydroxytamoxifen (OHT) to induce EMT for a total of 15 days. Every three days, cells were collected, lysed, and subjected to immunoblotting using the indicated antibodies. (**F**) GPX8 expression is upregulated in the TGFβ1 inducible system. A549 cells were treated with 5 ng/ml TGFβ1 to induce EMT for a total of 9 days. On each indicated day, cells were collected and subjected to immunoblotting using the indicated antibodies.

We then analyzed the cBioportal web-based resource tool (35, 36) to determine if there is any correlation pattern for GPX8 in patients. Specifically, we limited our analysis to breast cancer data generated by the cancer genome atlas (TCGA) (37). We first compared GPX8 expression relative to all the other genes and by Gene set enrichment analysis (GSEA) (38, 39) confirmed that in TCGA breast cancer samples GPX8 expression significantly correlates with the expression of known mesenchymal markers (Fig. S1A). Analysis of individual mesenchymal markers ZEB1, ZEB2, and CDH11 demonstrate a significantly high correlation with GPX8 (Pearson-0.57, 0.39, and 0.68, respectively), in contrast to the epithelial markers CDH1 (Pearson −0.17) (Fig. 1B). Together, both cell line and patient data confirm GPX8 expression correlates with the more aggressive, mesenchymal-like breast cancers.

Next, to validate the bioinformatics results, GPX8 expression between different cell lines generated from diverse cancer types was compared. In breast cancer cell lines GPX8 expression is higher in the more aggressive basal B subtypes (40) relative to luminal cell lines at both the RNA and protein levels (Fig. 1C, D). This expression pattern of GPX8 is similar to other known mesenchymal markers such as CDH11, ZEB1, VIM, and ZEB2, and is in contrast to the epithelial markers CDH1 and the epithelial cell junction protein occludin (OCLN) (41) (Fig. 1C, D). Additionally, in hepatocellular carcinoma cell lines (HCC) GPX8 expression is upregulated in the high-grade SNU-387 and SNU-423 (42) relatively to the low-grade cell lines HepG2 (Fig. 1D). Finally, in the highly metastatic melanoma cell line A375-MA2 (43), GPX8 expression is upregulated compared to the less aggressive parental cell line A375 (Fig. S1B). Together, these findings further demonstrated the upregulation of GPX8 in the high-grade tumors generated from different tumors.

Given the relatively high expression levels of GPX8 in mesenchymal-like cells, we then hypothesized that the EMT program regulates its expression. We found a gradual elevation in GPX8 expression in engineered human mammary epithelial (HMLE) cells induced to execute the EMT through the activation of Twist1 (8) (Fig. 1E). In parallel, we observed downregulation of the epithelial marker CDH1, confirming the activation of the EMT program. GPX8 expression elevation is also present in lung carcinoma cell line A549 treated with chronic transforming growth factor-beta 1 (TGFβ1) for nine days (44) (Fig. 1F). In these cells, GPX8 expression pattern correlates with mesenchymal markers CDH2, ZEB1, and vimentin (VIM) but anticorrelates with the epithelial marker CDH1 (Fig. 1F). Collectively, these results demonstrate that GPX8 expression is coupled with the expression of known mesenchymal markers, suggesting its possible role in tumor progression.

### GPX8 expression is associated with poor patient prognosis

The high correlation between GPX8 and EMT markers, predicts that GPX8 expression affects tumor aggressiveness. Indeed, the high expression of GPX8 in breast cancer samples correlates with the poor patient outcome as determined by the Kaplan-Meier Plotter tool (http://kmplot.com/analysis/) (45). Specifically, we analyzed the effect of GPX8 expression on breast cancer overall survival (OS), distance metastasis-free survival (DMFS), and relapse-free survival (RFS) (46). In all three parameters, GPX8 expression correlated with a significant reduction in the patient outcome (Fig. 2A). Moreover, the effect of GPX8 expression on patient outcome correlated with the aggressiveness state of the tumor samples. GPX8 expression levels in luminal A, the less aggressive type of breast cancer, did not demonstrate any apparent effect on patient outcome in contrast to the most significant effect observed in samples generated from basal breast cancer patients (Fig. 2A). Thus, further demonstrating its role in the more aggressive types of breast cancers. Correlation between high GPX8 expression and overall patient survival (OS) or first progression (FP) was also seen in samples from patients with lung (47) and gastric (48) cancers (Fig. 2B). Thus, expanding the role of GPX8 to other cancer types beside breast.

**Figure 2:**
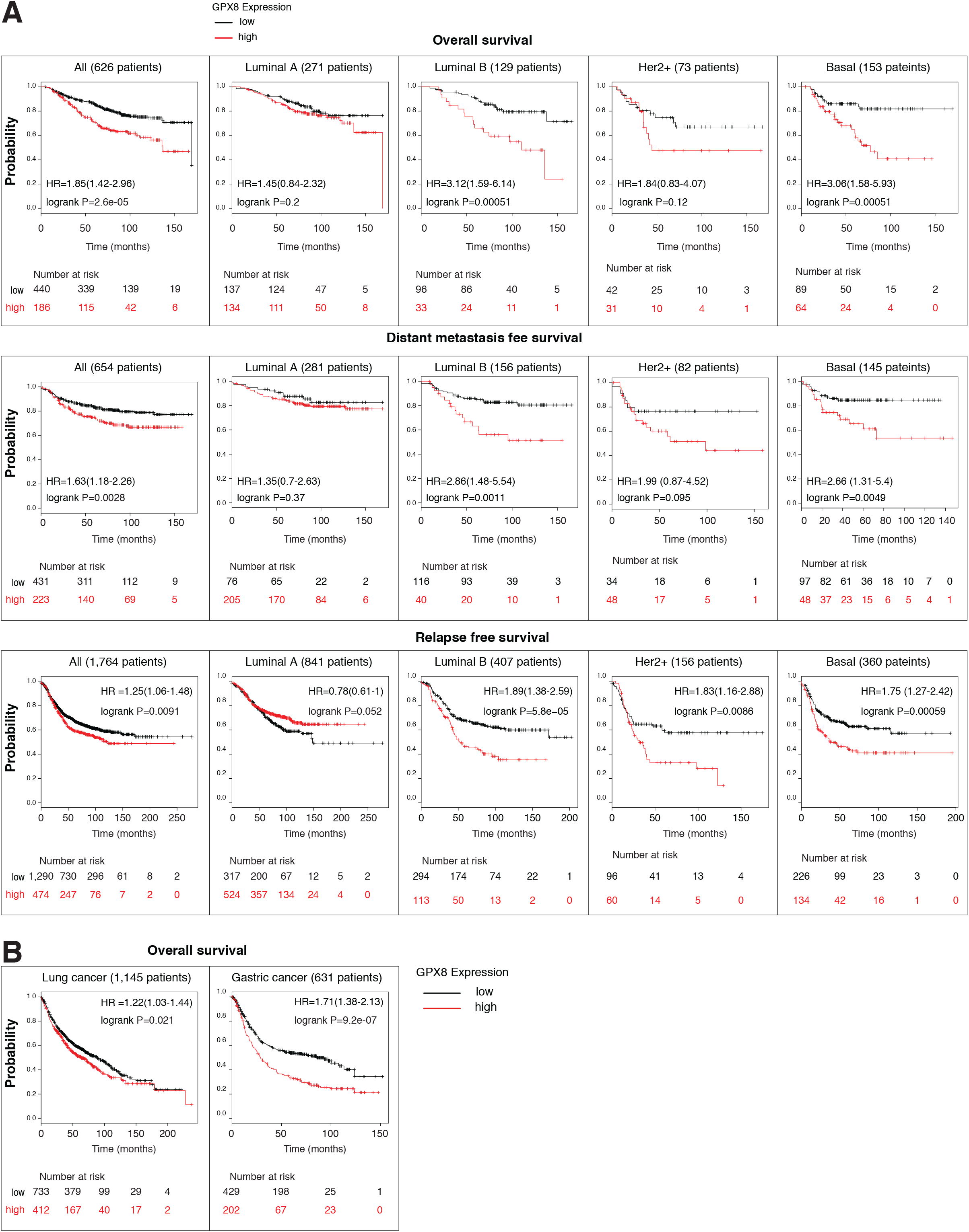
GPX8 expression is associated with poor patient prognosis: (**A**) Kaplan-Meier survival plots for patients with breast cancer divided into two groups (“high” and “low”) according to GPX8 expression. The columns represent data from all breast cancers (All), Luminal A, Luminal B, Her2+, and Basal. Each row represents a different type of patient survival plot, as indicated. These plots were generated in the Kaplan-Meier plotter website. The GPX8 228141_at Affymetrix ID symbol was used for all the analysis. The P value (P) and the number at risk were determined by the analysis tool. (**B**) The columns represent data from all lung cancers (Lung Cancer) and gastric cancers (Gastric Cancer).

Many of the aggressive tumor types express high levels of mesenchymal markers. However, not all of the mesenchymal markers are associated with poor patient prognosis. In order to demonstrate the important role of GPX8 in tumor aggressiveness, we compared its over-expression influence on the patient outcome with other mesenchymal markers (ZEB1, DPYD, and VIM). In contrast to GPX8 that resulted in a poor outcome, the other markers did not (Fig. S2), suggesting the active role of this enzyme on tumor aggressiveness. Together, GPX8 is upregulation in the more mesenchymal tumors, and its correlation with poor patient prognosis imply on its active role in the regulation of cancer cell aggressiveness.

### GPX8 silencing in TNBC MDA-MB-231 induces epithelial-like phenotype

The EMT-dependent elevation in GPX8 expression together with its correlation with poor patient outcome suggests it function as an EMT regulator. Therefore, we silenced GPX8 expression in the basal B breast cancer cell lines MDA-MB-231, using specific guides for Cas9-CRISPR system. GPX8 loss caused a significant alteration in cell morphology as they became smaller, rounder than WT cells, and clustered into island-like morphology in both GPX8-KO clone-1 and clone-2 (GPX8-KO-1, GPX8-KO-2, respectively, Fig. 3A), phenocopying epithelial cells characteristics. A phenotype that was abolished by the expression of WT-GPX8 in the KO background (Fig. 3A). This alternation in cell morphology was not associated with any changes in cell proliferation rate (Fig. S3A), suggesting that at least in MDA-MB-231 cells, this enzyme is not critical for proliferation.

**Figure 3:**
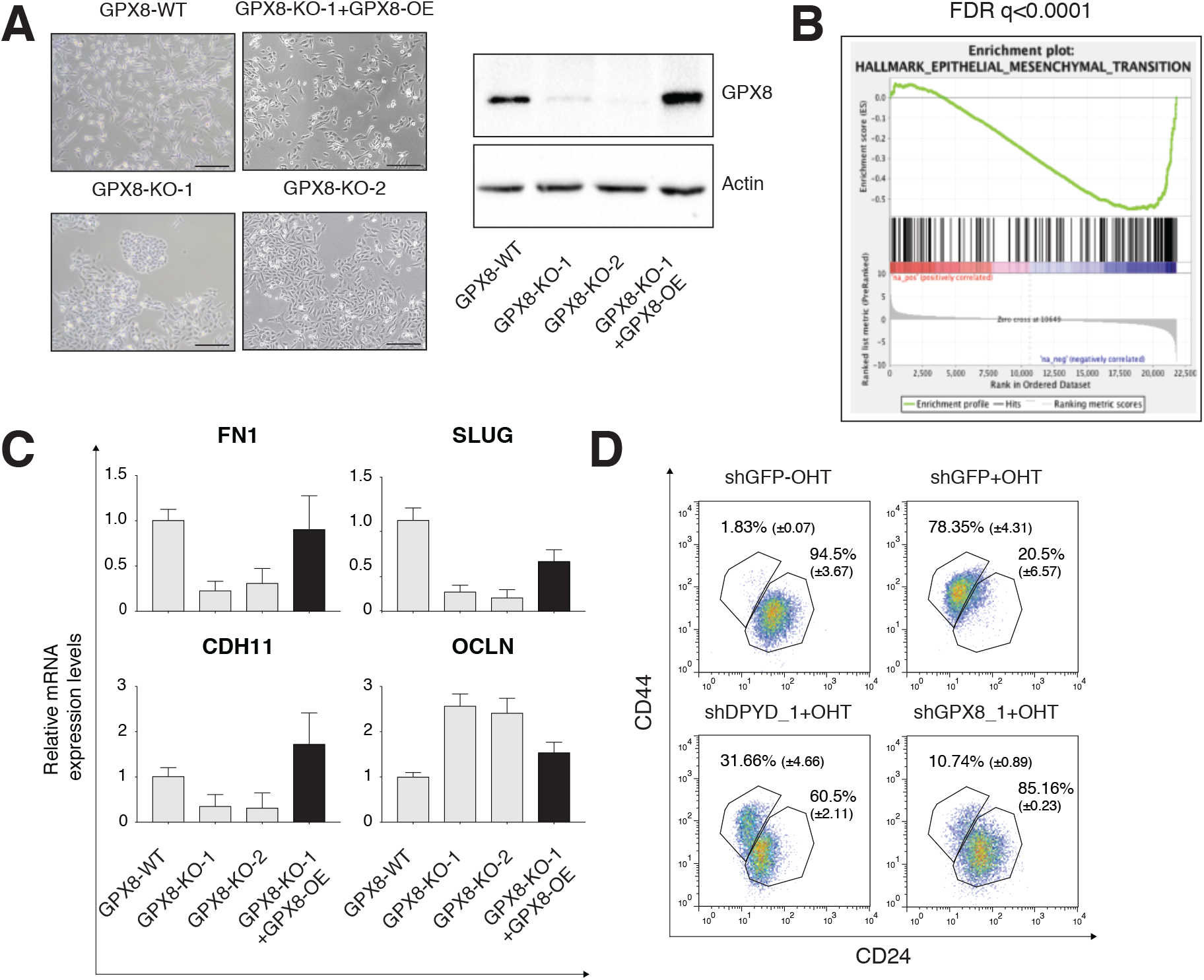
GPX8 loss results in epithelial-like characteristics: (**A**) KO of GPX8 in MDA-MB-231 cells induces epithelial-like morphology. GPX8 expression was attenuated in MDA-MB-231 cells using the CRISPR-Cas9 system. The cells were separated into single clones and for each one of them, GPX8 levels were measured by immunoblot using specific antibody against GPX8. GPX8-KO-1+GPX8-OE: GPX8 was reintroduced in the background of GPX8-KO-1 KO. In the right, the cells were lysed and subjected to immunoblotting with the indicated antibodies to demonstrate GPX8 levels in all samples. (**B**) GPX8 loss leads to reduced gene expression of EMT markers. MDA-MB-231 WT cells and GPX8-KO-1 were subjected to RNA-Seq analysis. The expression ratio between all genes (~22,000) was calculated and ranked based on the relative expression between the GPX8 WT and KO. The samples were then subject to gene-set enrichment analysis (GSEA). FDR q-value was computed by GSEA. (**C**) KO of GPX8 in MDA-MB-231 cells reduces the expression of known mesenchymal markers. The RNA was isolated from the different clones described above (a). The expression of the selected genes in the different colonies was determined by qRT-PCR. Each value represents the mean ± SD for n = 3. (**D**) GPX8 KO inhibits the EMT program. HMLE-Twist-ER cells were infected with the indicated hairpins. The cells were either left untreated or treated with OHT for 15 days, followed by FACS analysis of the cell-surface markers CD24 and CD44 to separate the epithelial and mesenchymal populations. The percentage of cells in each gate is presented.

We then subjected WT cells and GPX8-KO-1 (clone-1) to RNA-Seq analysis to systematically identify the molecular mechanisms inducing these cellular changes. GSEA confirmed that GPX8 loss caused a significant reduction in “the hallmarks of the epithelial-mesenchymal transition” (FDR q-value<0.0001; Fig. 3B) gene set. Specifically, GPX8-KO results in the downregulation of the known mesenchymal markers such as FN1, SLUG, and CDH11 as well as upregulation of the epithelial marker OCLN (2) (Fig. 3C).

The role of GPX8 in the maintenance of the mesenchymal state urged us to further characterize its role in the EMT program. To this end, we knocked down GPX8 expression in HMLE-Twist-ER cells using specific short hairpins RNA (shRNA). GPX8 reduction caused changes in the EMT-associated cell surface expression, namely CD24high/CD44low to CD24low/CD44high (Fig. 3D). Additionally, GPX8 expression is essential for the Twist-dependent upregulation of ZEB1 and CDH2 and the repression of CDH1 (Fig. S3B). Collectively, these data suggest that GPX8 expression plays an essential role in maintaining the cell’s mesenchymal properties by regulating the expression of particular EMT genes.

### GPX8 plays a role in the aggressive characteristics of MDA-MB-231 cells

Upon the execution of the EMT program, the carcinomas gain mesenchymal-like characteristics such as increased migration capability (1). Since GPX8 loss induces epithelial-like gene expression profile, we next asked whether it also affects the mesenchymal features of these cells. To this end, we performed wound healing and migration assays on the different KO clones. The Incucyte Live-Cell Analysis System was used to monitor the migrating cell in real-time. We observed that GPX8 loss significantly reduced wound healing efficiency that was rescued in the re-expression of GPX8 in the GPX8-KO background (Fig. 4A and Fig. S3A). In addition, the cells demonstrated a significant reduction in their motility capability in the transwell migration assay (Fig. 4B). Together, these experiments demonstrate GPX8 to have a vital role in MDA-MB-231 mesenchymal features maintenance, such as the ability to migrate.

**Figure 4:**
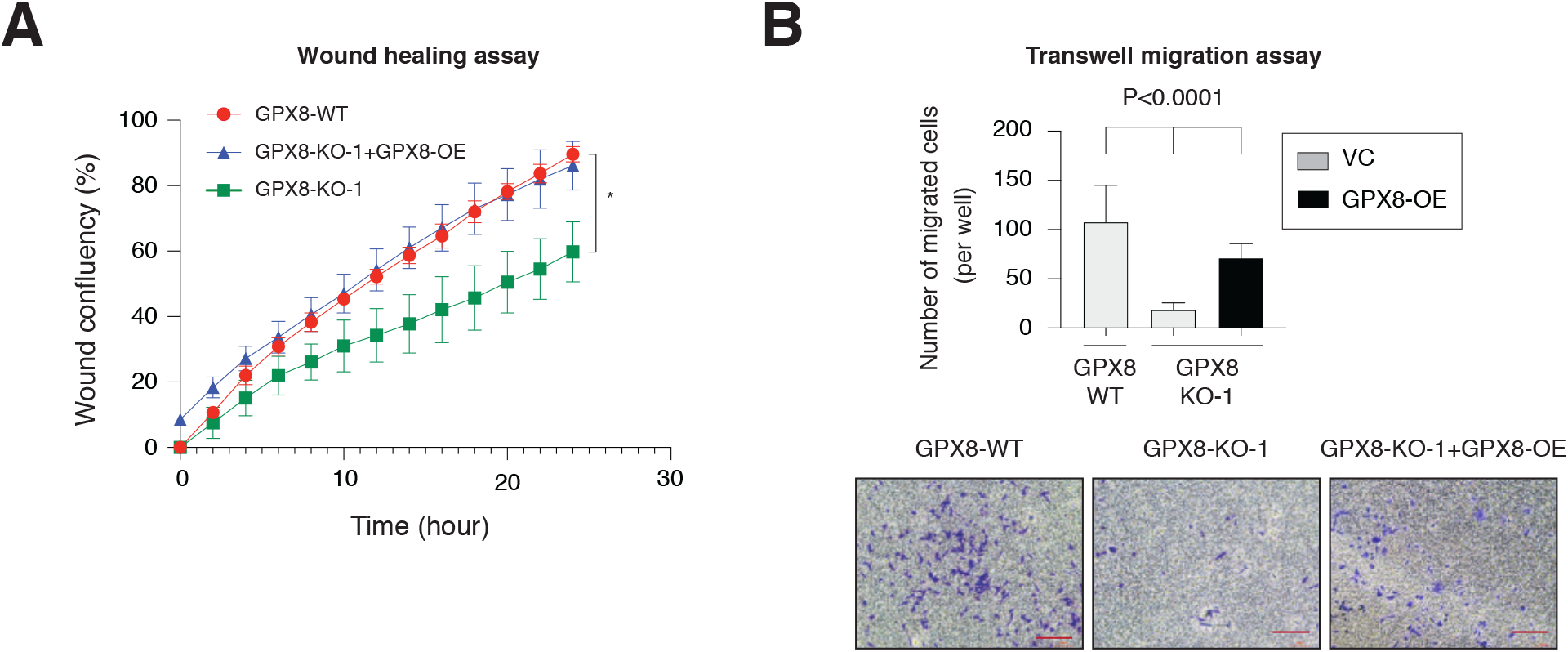
GPX8 loss inhibits cell migration in the MDA-MB-231 breast cancer cell line. (**A**) (Top panel) Quantification of wound confluence of the indicated GPX8 KO clones during 24 hours. For each sample n=3, *-P=0.01 that was determined by T-test using Prism software. (**B**) GPX8 loss inhibits the MDA-MB-231 migration capabilities. (top panel) The migration capability of the different samples was determined in a transwell assay. The data are reported as the number of migrated cells per 10,000 seeded cells; each value represents the mean ± SD for n = 3. P value was determined by Student T-test using Prism software (low panel). Representative cell migration images of each sample. Bar = 200μm.

### GPX8 regulates the stemness properties of cancer cells

Previously we performed a FACS-based screen in HMLE-Twist-ER cells to comprehensively identify the MMS genes playing essential roles in the EMT program (10). Specifically, we compared the expression levels of the epithelial CD24 marker and the mesenchymal/stem cell marker CD44 (8, 49). This screen, resulted in a list of 16 hits genes, among them GPX8, which we confirmed to be essential to the EMT program (Fig. 3). Since GPX8 regulates stem cell marker CD44, we asked whether GPX8 regulates the cancer stemness capabilities. Several of the basal B breast cancer cell lines, including MDA-MB-231, have stem-like properties (50) as they express specific markers and can initiate tumors in mice. We revealed that GPX8 loss resulted in the shift of cell-surface markers from mesenchymal/stemness (CD24^low^, CD44^high^) to an epithelial-like (CD24^high^, CD44^low^) profile (8, 49) (Fig. 5A). In addition, in GPX8 KO cells the expression of known stemness markers such as integrin-β4 (ITGB4) (51) (Fig. 5B), and nerve growth factor receptor (NGFR, p75NTR (52)) was suppressed (Fig. 5C).

**Figure 5:**
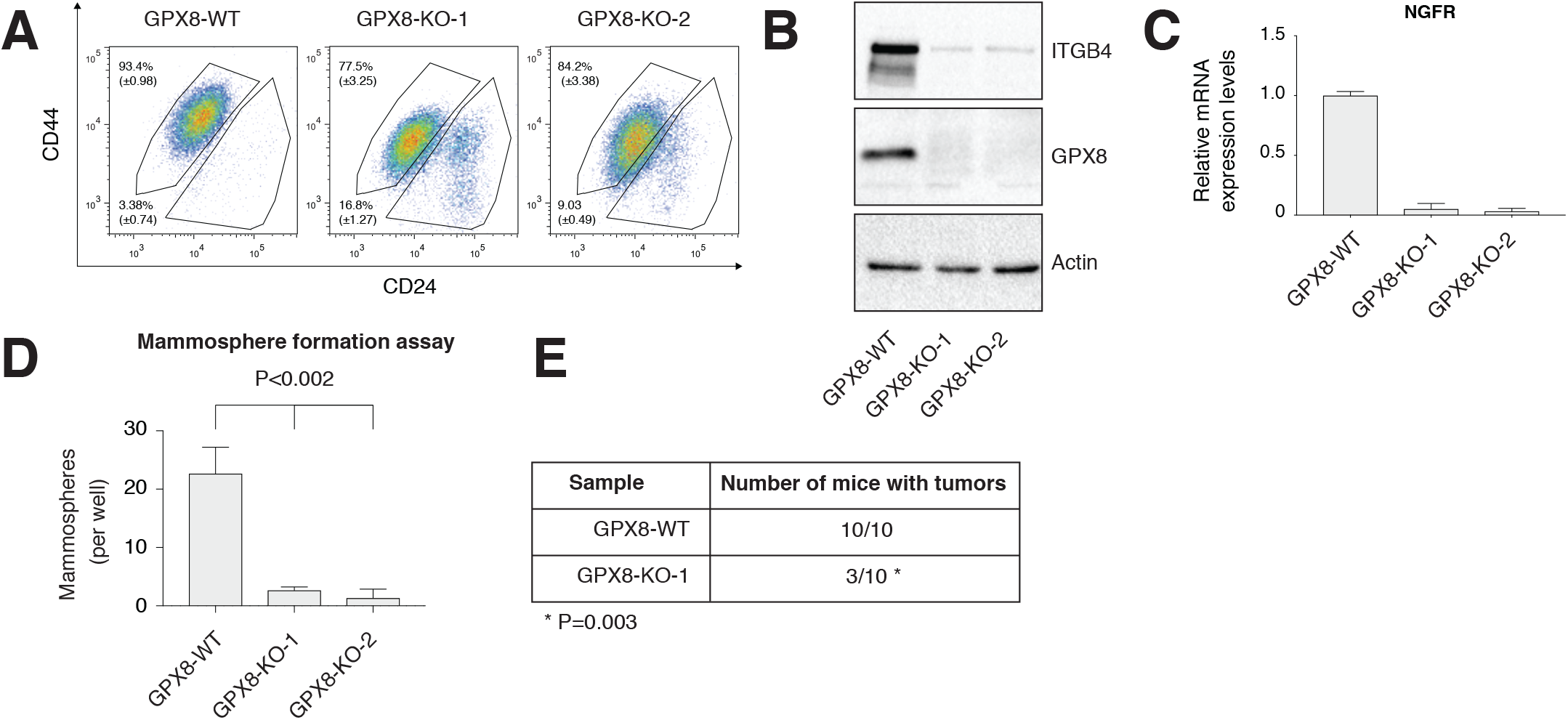
GPX8 loss in MDA-MB-231 cell affects cancer stemness. (**A**) GPX8 KO results in a CD24^hi^/CD44^low^ population. The GPX8 clones as in Figure 3A were subjected to FACS analysis of the cell-surface markers CD24 and CD44 to separate the epithelial and mesenchymal populations. The percentage of cells in each gate is presented. (**B**) GPX8 expression levels correlate with the stem cell markers. Cells from the different GPX8-KO clones were subjected to immunoblotting with the indicated antibodies. (**C**) GPX8 expression levels correlate with the stem cell marker NGFR. The RNA from the different GPX8-KO clones was isolated and the level of NGFR was determined using specific primers. (**D**) GPX8 expression in MDA-MD-231 cells correlates with the cells’ ability to form mammospheres. Quantification of *in vitro* mammosphere formation by cells from the different clones was performed. The data are reported as the number of mammospheres formed per 600 seeded cells; each value represents the mean ± SD for n = 3. P value was determined by Student T-test using Prism software. (**E**) GPX8 loss affects tumor formation in mice. Female NOD-SCID mice were injected with 10^6^ cells generated from the different clones. After six weeks, the proportion of animals bearing tumors was assessed and presented.

For an *in vitro* stemness functional assay, we measured the capability of the GPX8-KO clones to form mammospheres. We found that GPX8 KO formed very poorly mammospheres (Fig. 5D and Fig. S4A) in comparison to the WT, and KO-rescue cells. We then determine the effect of GPX8 loss on the ability of the MDA-MB-231 to form tumors in female NOD-SCID mice. Specifically, we injected the same number of cells originated from WT and GPX8-KO-1 cells into the fat pad of female mice, and after six weeks we measured the number and size of the generated tumors. While all the injected mice with the WT cells generated tumors, only three mice were positive for the tumors with GPX8-KO-1 (Fig. 5E) with one of the tumors only 0.05gr (Fig. S4B). Together, these results suggest that GPX8 cellular function is to maintain the cancer cells stemness characteristics.

### GPX8 regulates the secretion of autocrine cytokines

Following the evidence supporting that GPX8 expression plays a critical role in cancer stemness, we seek to understand the underlying mechanism. The localization of GPX8 in the ER and its contribution to disulfide bond regulation (24) suggest its role in the maturation of secreted factors such as cytokines. The fact that our RNA-seq GSEA analysis revealed a GPX8-dependent expression of the inflammatory response (“Hallmark Inflammation Response” (Fig. 6A)) led to further support to this possibility. We, therefore, assayed to determine whether the knockout of GPX8 affects the production of secreted factors. To this end, we incubated GPX8-KO-1 cells with WT conditioned medium and found upregulation of the EMT marker FN1, and the stem cell markers CD44 and ITGB4 (Figure 6B). Therefore, validating the role of GPX8 as a regulator of EMT/stemness-associated secreted factors.

**Figure 6:**
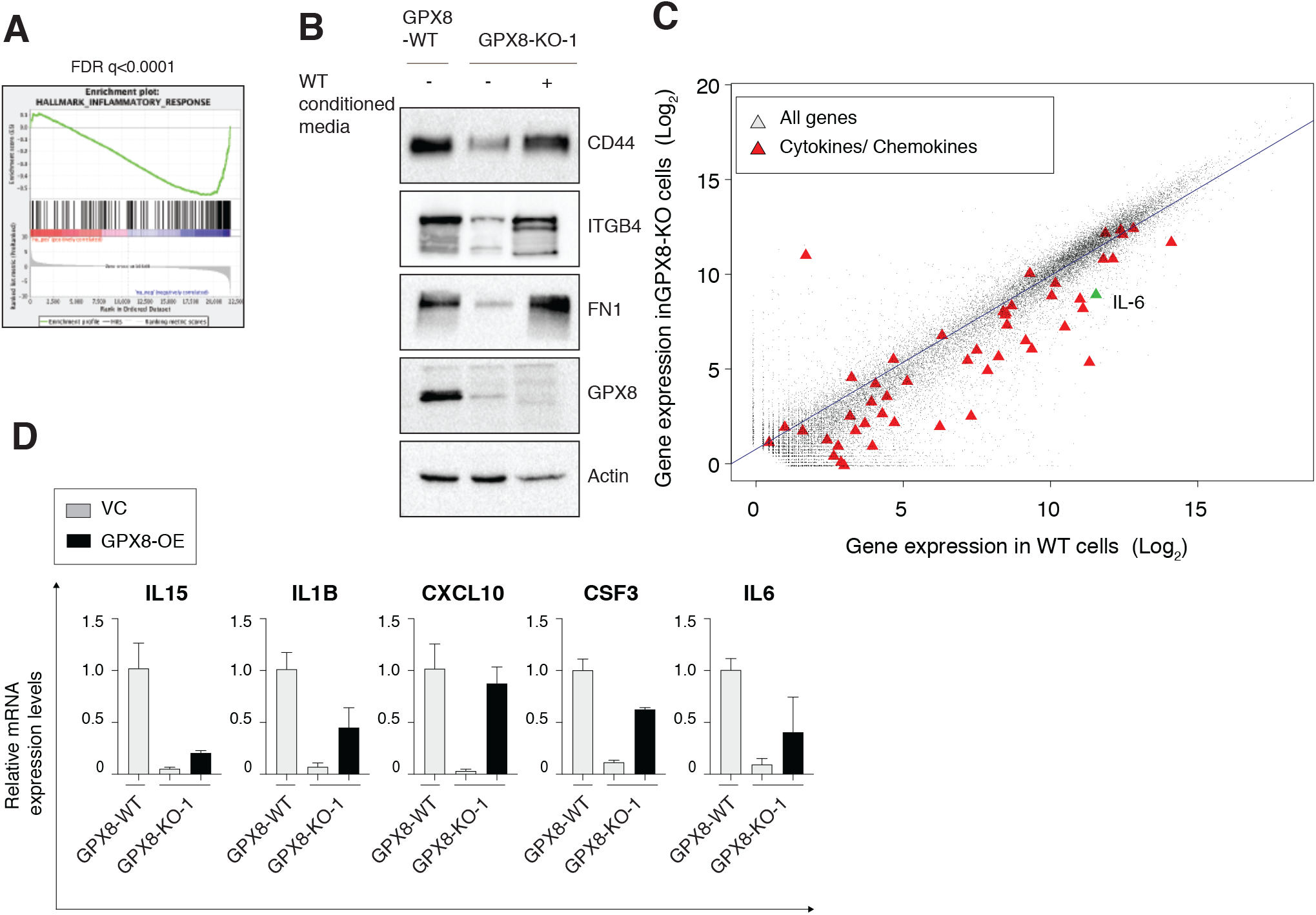
GPX8 KO impairs cytokine production in MDA-MB-231 cells. (**A**) GPX8 loss leads to a reduction in the gene expression pattern for “Hallmark Inflammatory Response”. MDA-MB-231 WT cells and GPX8-KO-1 were subjected to RNA-Seq analysis. The expression ratio of all genes was calculated and ranked based on the relative expression in GPX8 WT and KO. The samples were then subjected to gene-set enrichment analysis (GSEA). The FDR q-value was computed by GSEA. (**B**) Conditioned media from WT cells can rescue the expression of EMT markers. Conditioned media form the WT was added to GPX8-KO-1cells for three days. The cells were then lysed and subject to immunoblot using the indicated Abs. (**C**) GPX8 loss induces a global reduction in cytokine expression. The expression level in WT and GPX8 KO cells is presented in a scatter plot. Cytokines and chemokines are presented as a red triangle apart from IL-6 expression which is presented in green. (**D**). KO of GPX8 in MDA-MB-231 cells reduces the expression of selected cytokines. The RNA was isolated from WT and GPX8-KO-1 cells. The expression of the selected genes was determined by qRT-PCR. The expression of each gene was expressed as a ratio of its expression in WT (black) and KO (gray) cells. Each value represents the mean ± SD for n = 3.

To systematically identify these secreted factors, we generated a list of all the predicted secreted proteins using the “Human Protein Atlas” database (https://www.proteinatlas.org). From the 2249 genes listed, we chose 78 genes encoding for cytokines and 44 for chemokines. To identify the secreted factors in the growth media systematically, we analyzed our RNA-seq results to identify the cytokines and chemokines that are expressed in these cells, resulting in a restricted list of only 50 genes (cytokines gene set, (CGS)) (supplement Table-1). Analysis of the gene expression profile identified the CGS to be significantly reduced in the GPX8 KO cell relative to the WT (Fig. 6C and Fig. S5A). This large-scale analysis was then individually validated by examining the expression of selected cytokines and confirmed their reduction in GPX8-KO cells, and their upregulation in GPX8 overexpression in KO background (Fig. 6D). We found that GPX8 loss reduced the expression of interleukine-15 (IL-15), interleukine-1β (IL1B), C-X-C motif chemokine ligand 10 (CXCL10), colony stimulating factor 3 (CSF3), and interleukine-6 (IL-6). Together, we identify GPX8 as a global regulator for the production of fundamental cancer-associated cytokines.

### GPX8 regulates the IL6-STAT3 signaling pathway

Activation of the IL-6-Signal Transducer And Activator Of Transcription 3 (STAT3) signaling pathway is associated with tumor progression, the EMT program (53), and provides a critical link between inflammation and cancer (54). Analysis of the MERAV database supports IL-6 role in aggressive cancers as its expression is significantly upregulated in cell lines baring mesenchymal markers relative to epithelial cells (Fig. S5B), consistence with studies that identified expression elevation in basal-like than luminal cell lines (55). IL-6 functions in a feedback loop, where it interacts with the IL6 receptor (IL6R) and activate the Janus kinase (JAK)-STAT3 signaling pathway (56). JAK-STAT3 activation induces the expression of many cancer-associated factors among them is the IL-6 itself (56). This activation of IL-6-JAK-STAT3 signaling pathway has an essential role in regulating the equilibrium toward CSCs, inducing the upregulation of stemness markers such as CD44 (5, 57). GSEA analysis of our RNA-seq results (as described above) indicated that loss of GPX8 caused a significant downregulation of “Hallmark of IL-6-JAK-STAT3 signaling” (Fig. 7A), suggesting GPX8 as a key regulator of this signaling pathway.

**Figure 7:**
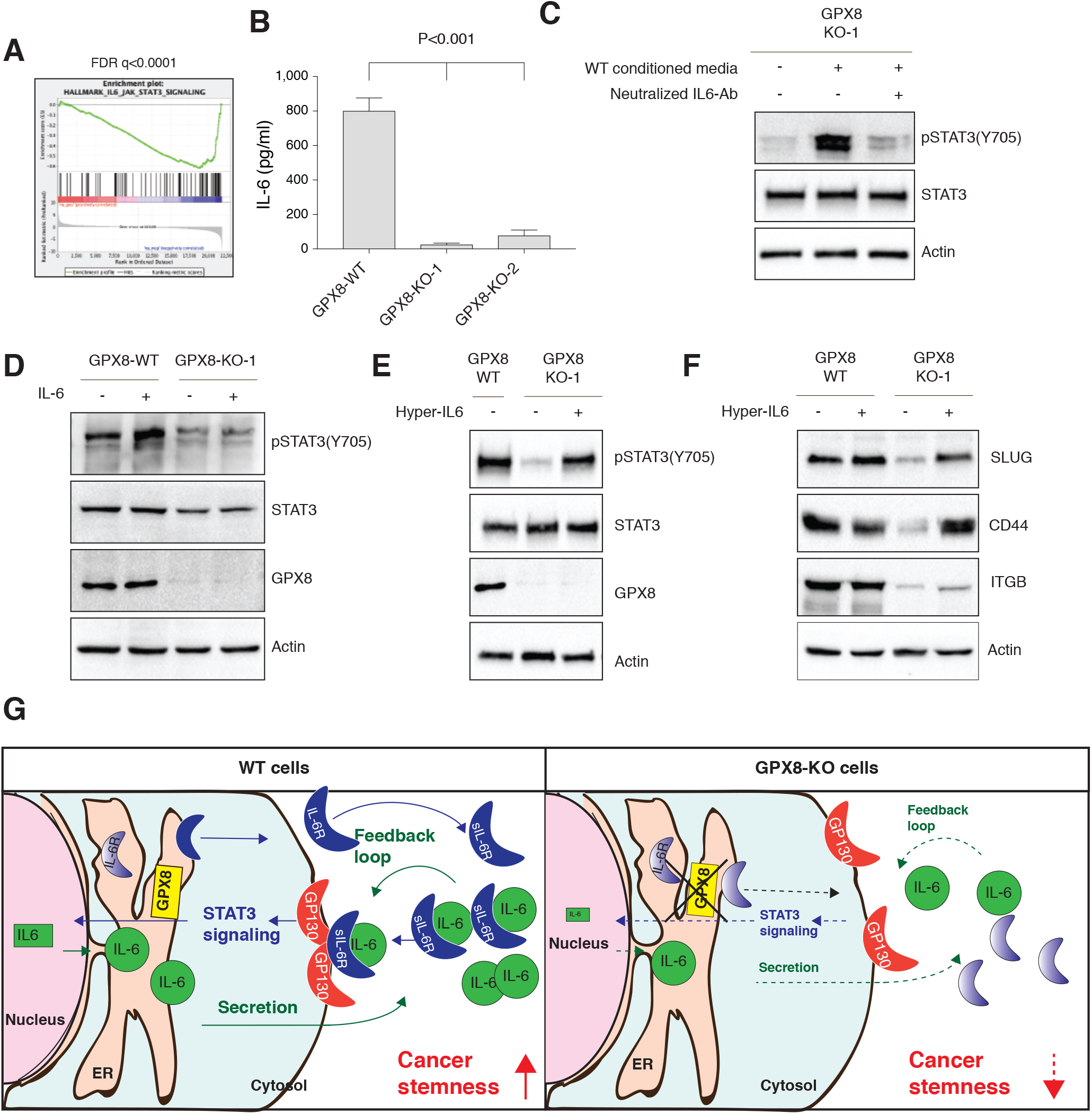
GPX8-KO cellular effect are mediated by IL-6-JAK-STAT3 signaling pathway. (**A**) GPX8 loss leads to a reduction in the gene expression pattern for “Hallmark IL-6 JAK STAT3 signaling”. MDA-MB-231 WT cells and GPX8-KO-1 were subjected to RNA-Seq analysis. The expression ratio of all genes was calculated and ranked based on the relative expression in GPX8 WT and KO. The samples were then subjected to gene-set enrichment analysis (GSEA). The FDR q-value was computed by GSEA. (**B**) IL-6 level is reduced in GPX8 KO cells growth media. Cells growth media was collected from each of the indicated samples for two days. The level of IL-6 was determined using a specific ELISA kit each sample (n=18). P value was determined by Student T-test using Prism software. (**C**) Conditioned media from WT cells induces STAT3 tyrosine phosphorylation (Y705) via IL-6. Conditioned media form the WT was added to GPX8-KO-1cells for one hour. The cells were then lysed and subject to immunoblot using the indicated Abs. STAT3 signaling is reduced by the addition of neutralized IL-6 antibodies to the conditioned media. (**D**) GPX8-KO cells do not respond to IL-6 treatment. WT and GPX8 KO cells were stimulated with 5 ng/ml IL-6 for 3 days. Cells were subjected to immunoblot using the indicated Abs. (**E**) Hyper-IL6 rescues the STAT3 signaling pathway in GPX8-KO cells. WT and GPX8-KO cells were stimulated with hyper IL-6 (50ng/ml) for 45 min. The cells were then lysed and subject to immunoblot using the indicated Abs. (**F**) Hyper-IL6 induces EMT markers expression in GPX8-KO cells. WT and GPX8 KO cells were stimulated with hyper IL-6 (50ng/ml) for 3 days. The cells were then lysed and subject to immunoblot using the indicated Abs. (**G**) A scheme representing the role of GPX8 in IL-6 maturation and IL-6 receptor activation. Green arrows represent the IL-6 secretion, dark blue arrows represent intracellular IL-6 signaling. Dashed arrows and reduced font size represent impaired secretion and intracellular signaling. In WT cells, IL-6 and IL6R are processed in the ER gradient color represented proteins before maturation and solid color after maturation.

We next set out to validate GPX8 function as a significant regulator of the IL6-JAK-STAT3 signaling cascade. We found by ELISA analysis, a significant reduction in IL-6 secretion in GPX8-KO cells by about 90-95% compared to WT cells (24.5-77.42pg/ml vs. 800.0pg/ml, respectively) (Fig. 7B). In addition, neutralizing IL-6 antibodies abolished STAT3 phosphorylation upon treatment with conditioned media generated from WT cells (Fig. 7C). Together, indicating that GPX8 is a key regulator of IL-6 secretion, which functions as JAK-STAT3 signaling pathway activator.

We noticed that GPX8 affect IL6-JAK-STAT3 feedback loop; however, the specific component of this signaling pathway regulated by GPX8 is still unclear. Curiously, we found that treatment of GPX8-KO cells with recombinant IL-6 failed to activate STAT3 (Fig. 7D), despite the presence of GP130 on these cells (Fig. S5C). Interestingly, assessment by FACS analysis indicated that IL6R expression levels on the surface of MDA-MB-231 GPX8-WT, GPX8-KO, and GPX8-KO rescue cells were relatively undetectable (Fig. S5D). Indicating, that the “IL-6 classic signaling”, which is composed of secreted IL-6 and transmembrane IL6R does not apply in these cells (58). Therefore, these observations suggested that IL-6 signaling induced by GPX8-WT conditioned media (Fig. 7C) is mediated by an alternative mechanism designated as “IL-6 trans-signaling”. In this mechanism GP130 activation is mediated via a complex of IL-6 together with its soluble IL-6R (sIL-6R), as described previously (58). Thus, incubation of GPX8-KO cells with a recombinant IL-6/sIL-6R fusion protein, called Hyper-IL-6 (59), overcome the absence of IL-6R expression and activate the IL-6 signaling cascade in these cells (Fig. 7E). Moreover, treatment with Hyper-IL-6, also rescued the expression of the EMT markers SLUG and CD44 in the GPX8-KO cells (Fig. 7F). Thus, GPX8 regulates STAT3 signaling via the production and secretion of sIL-6R, which activates the EMT program through IL-6 trans-signaling mechanism.

## Discussion

We identify a novel GPX8-IL-6 axis which activation during the EMT program promotes the acquisition of stemness characteristics in breast cancer cells. GPX8 expression correlates in patients with known mesenchymal markers and is associated with poor patient prognosis. Moreover, the EMT program directly upregulates GPX8 expression, which is essential for the maintenance of cancer cells’ aggressive phenotype. GPX8 induces these cellular alternations through the regulation of IL-6-STAT3 signaling cascade. A signaling pathway that plays an essential role in promoting breast cancer aggressiveness (56). In addition, the activation of this STAT3 signaling pathway induce the expression of the stemness marker CD44 in hepatocellular carcinomas (60). Specifically, we demonstrate GPX8 regulation on IL6R that fails to activate the IL-6 trans-signaling. Thus, we propose a model by which GPX8 regulates the proper production of IL6R through the interaction with IL-6 activated the JAK-STAT3 signaling. This signaling cascade induces the expression of IL-6 that function as an autocrine factor which induces the EMT program (Fig. 7G). GPX8 loss impairs the proper production of IL6R, leading to a reduction in the JAK-STAT3 signaling pathway activation, resulting in a decreased yield of cytokines production, which includes IL-6. This reduction in IL-6 levels results in the cellular loss of their aggressive phenotype (Fig. 7G).

Our model indicates that GPX8 plays a central role in the EMT program. Two EMT-induction system models (TGFβ and Twist) upregulates GPX8 expression (Fig. 1), implying its critical role in this program. A recent study identified two hypoxia-response elements within GPX8 promotor, demonstrating HIF1α as its primary expression regulator (30). HIF1α plays a critical regulatory role in the EMT program (61) by repressing E-cadherin expression (62) or by directly activating the expression of the key EMT transcription factors TWIST (63), ZEB1 and ZEB2 (2). Additionally, HIF1α promotes triple negative breast cancer tumorigenesis (64). Thus we predict that this can be one possible mechanism regulating GPX8 expression during EMT.

The EMT program was first reported in embryonic development, where selected cells change their epithelial identity and gain mesenchymal-like characteristics (2). Further studies expanded the role of the EMT program to be included in wound healing, fibrosis, and cancer. Fibrosis is a complex disease associated with reduced organ function by the excessive synthesis and accumulation of extracellular matrices due to the accumulation of fibroblasts and myofibroblasts (65). Multiple causes such as physical injury and virus infection trigger fibrosis (66) by cellular mechanisms which are reminiscent of oncogenesis. Interestingly, ROS (66) and selected cytokines such as IL-6 (67) drive the activation of fibroblast during fibrosis. This similarity in molecular mechanism between fibrosis and cancer-dependent EMT suggests that GPX8 is very likely to play a role in fibrogenesis as well. Thus, we predict that GPX8 function is not limited to type 3 of EMT (association with cancer progression (68)) and can play a significant role in the other types of EMT (during embryogenesis and fibrosis). Therefore, GPX8 is a suitable candidate for drug target development aim to inhibit both cancer aggressiveness and fibrosis.

The presence of the unconventional amino acid, selenocysteine in their activity site divides the GPx family of enzymes into two groups, GPX1-4, and GPX6 that do contain, and GPX5, GPX7-8 that do not. Unambiguous phylogeny analysis of the GPX family further subdivided the eight members into three groups, whereby GPX4, GPX7, and GPX8 belong to the same evolutionary branch (69). GPX4 was found to be one of the primary regulators of ferroptosis (70), a form of apoptotic cell death (71). A selective GPX4 inhibitor, RSL3, induces cell death in epithelial cancer-derived cell lines expressing mesenchymal markers such as ZEB1 (72). In addition, persister cancer cells, which are resistance to lapatinib treatment and upregulate mesenchymal and stem cell markers, are vulnerable to GPX4 inhibition by RSL3 (73). This mutual mesenchymal cell-dependent function of GPX4 and GPX8 suggest a conserved role for this sub-family of GPx in aggressive cancer cells.

As part of their production, cytokines such as IL-6 and their receptor (IL6R) are processed in the ER where they acquire their proper folding mediated by disulfide bonds (74, 75). Specifically, the extracellular soluble domain of IL6R, which interacts with the ligand contains four conserved cysteines that form disulfide bonds (76). The ER localization of GPX8 and its functions as a regulator of proteins disulfide bonds formation (24) suggest that it regulates IL6R proper folding. Here we identified that GPX8 functions as a critical regulator of cytokine production in aggressive cancers, suggesting for a robust cellular mechanism mediated by this enzyme. Thus, we suggest a model by the EMT program elevates GPX8 expression to function as a regulator of the proper production of cytokines and their receptors. These fully functional autocrine cytokines are secreted and induce the stemness properties in these tumors via the JAK-STAT3 signaling cascade (56). However, In GPX8-KO cells, both the cytokines and IL6R folding is impaired, thus braking this feedback loop, which shifts the equilibrium toward the less aggressive phenotype (Fig. 7F).

## Acknowledgements

We thank the members of the Y.D.S. laboratory. This work was supported by the Israel Science Foundation (Grant 1816/16) and the Israeli Cancer Association (Grant 20180062) and the Hebrew-University start-up funds. S.B. is supported by the Lady Davis Fellowship for post-doctoral researchers at The Hebrew University of Jerusalem.

## Author contributions

A.K. and Y.D.S. designed the study. A.K. performed most of the experiments except for the bioinformatics. S.B. performed most of the functional assay and in vivo studies. M.B.Y together with A.A.R performed many of the western blots and qPCR studies. M.S performed the Il-6 ELISA and TGFB experiments. M.L. did many of the cloning and contribute to experimental design. J.H.A provides the hyper IL-6 reagents and contribute to experimental design. Y.D.S and M.L wrote the paper with input from all the authors.

## Methods

### Cell Lines and Cell Culture

The immortalized human mammary epithelial cells expressing OHT-inducible Twist (HMLE-Twist-ER), and Naturally Arising MEsenchymal Cells (NAMECs) have been described before (8, 77). HMLE-Twist-ER and NAMEC cells were maintained in MEGM (Lonza) growth media. The cell lines A549, EVSA-T, BT-474, MCF7, MDA-MB-231, MDA-MB-436, Hs-578-T, HepG2, SNU-387, and SNU-423 were obtained from ATCC and were maintained in DMEM supplemented with 10% FBS. All cells were cultured at 37°C with 5% CO2. For EMT induction in HMLE-Twist-ER, cells were treated with 4-hydroxytamoxifen (OHT) (Sigma-Aldrich, H7904) at a final concentration of 10nM for the indicated number of days. For EMT induction in A549, cells were grown in 5% FBS for 48 h and then treated with 5 ng/ml TGFβ (Peprotech, New Jersey, USA) for the number of indicated days (44).

### Antibodies

Antibodies were obtained from the following sources: GPX8 (HPA036720) from Sigma Aldrich, CDH1 (3195), CDH2 (13116), Actin (4970)\(3700), ZEB1 (3396), VIM (5741), ITGB4 (14803), CD44 (for immunoblotting (3570)), p-sSTAT3-Tyr705 (9145), STAT3 (9139), from Cell Signaling Technology; FITC-labeled anti-CD24 (555427), APC-labeled anti-CD44 (559942) from BD Bioscience; IL6-R anti mouse (sc-373708) HRP-labeled anti-mouse and anti-rabbit secondary antibodies from Santa Cruz Biotechnology. GP130 anti rabbit (06-291) from Upstate. For IL-6 neutralization assay we used goat Anti-Human IL-6 (500-P26G), (Peprotech).

### Cell proliferation assay

Cells were seeded in white 96-well plates (Greiner) at a density of 500 cells/well (MDA-MB-231 WT, GPX8 KO clone 1 and 2). Cell viability was assessed with Cell Titer-Glo (Promega) at day 0, 2, and 4 days following seeding and luminescence measured with a Cytation 3 (Biotek).

### Cancer sample analysis

Kaplan-Meyer analysis of the data from breast and other cancer samples were analyzed and generated by the Kaplan-Meier Plotter website (http://kmplot.com/analysis/) (45). GPX8 was search as a gene symbol and the 228141_at Affymetrix ID was chosen. The Auto select best cutoff was selected as well. The obtained KM plots are presented. The statistics is generated by the website itself.

### CRISPR/Cas9-mediating and shRNA generation of knockout/knockdown cell lines

We used the CRISPR-Cas9 medicated genome editing to achieved gene knockout, using pLentiCRISPR v1 (Addgene Plasmid #70662) in which the sgRNA and Cas9 are delivered on a single plasmid. Editing of the GPX8 locus in MDA-MB-231 cells was accomplished by infecting cells with the “pLentiCRISPR” plasmid into which an sgRNA targeting the GPX8 locus had been cloned. Infected cells were subjected to single cell cloning by limiting dilution in 96-well plates. Editing of the GPX8 locus was confirmed by Sanger sequencing of the targeted locus. The following sense (S) and antisense (AS) oligo- nucleotides were cloned into pLentiCRISPRv1:

sgGPX8 (S): caccgAACTGCAAATACCTTTGCTC
sgGPX8 (AS): aaacGAGCAAAGGTATTTGCAGTTc

**shRNA**- pLKO.1 lentiviral plasmid encoding shRNAs targeting GPX8 (TRCN0000140309), DPYD (TRCN0000221382), and GFP (TRCN0000072178) were assembled into viruses. HMLE-Twist-ER cell were infected with the corresponding plasmids and then selected using puromycin.

### Statistical Analysis

All statistical analyses were performed using the Prism 8.0 statistical analysis program (GraphPad), and by the R software. If not indicated otherwise, all the p values in the figures measured between the indicated samples were quantified using Student’s t test.

