## Supplemental information for "Glutathione Peroxidase 8 (GPX8)-IL6 axis is essential in maintaining breast cancer mesenchymal stem-like state and aggressive phenotype"

#### *Supplementary Information*

##### *Cell Lysis and Immunoblotting*

Cells were rinsed once with ice-cold PBS and lysed with RIPA lysis buffer (20mM Tris [pH 7.4], 137mM NaCl, 10% glycerol (vol/vol), 1% Triton X-100, 0.5% (wt/vol) deoxycholate, 0.1% (wt/vol) SDS, 2.0mM EDTA [pH 8.0] and one tablet of EDTA-free protease inhibitor (Roche) and phosphatase inhibitor cocktail mixes A and B (100X) (Bimake, USA). The protein concentration was determined by Bradford (BioRad, 500-0006). The supernatants were then separated by a 10% SDS-PAGE, transferred onto a 0.45µ PVDF membrane (Merck 0.45µ) and probed with the appropriate antibodies.

##### *Virus Production*

HEK-293T cells were co-transfected with the pLentiCRISPR sgRNA library, the VSV-G envelope plasmid and the ΔVPR lentiviral packaging plasmid and retroviral packaging plasmids Gag-Pol and VSV-G, using X-TremeGene 9 Transfection Reagent. The virus-containing supernatant was collected 48 h after transfection and spun for 5 min at 400 g to eliminate cells.

##### *Animal Studies*

MDA-MB-231 WT cells and GPX8-KO-1 were injected into the fat pad of female NOD-SCID mice (1,000,000 cells per mouse). After six weeks the tumors were removed and weighed.

##### *FACS Analysis*

For FACS analysis, cells were prepared according to standard protocols and suspended in 1% Serum/PBS on ice prior to FACS. Cells were sorted on a BD Accuri C6 or analyzed FlowJo software (Tree Star, Ashland, OR).

##### *RNA Preparation and RT-PCR Analysis*

Total RNA was isolated from cells using the NucleoSpin® RNA Kit (MACHEREY-NAGEL, Germany) and reverse-transcription was performed using qScript cDNA Synthesis Kit (Quantbio, USA). The resulting cDNA was diluted in DNase-free water (1:10) before quantification by real-time quantitative PCR. The mRNA transcription levels were measured using SYBR Green PCR master mix Blue Mix HI-ROX (PCR Biosystems, London, UK) and StepOnePlus (Applied Biosystems, CA, USA). All data are expressed as the ratio between the expression level of the target gene mRNA and that for actin. Primers used for qRT-PCR were obtained from Integrated DNA Technology and are listed in**:** List of cytokines used for the RNA-seq analysis are list in supplement Table-2.

***Mammosphere formation Assay***

Six hundred cells were seeded per well in 96-well plates after coated with growth-factor-reduced Matrigel (cat no. 356230; Corning). Plates were kept growing for 6-9 days under observation and every two days once 2% of new growth-factor-reduced Matrigel was added. Sphere were counted manually under the microscope and scored as a percentage of spheres, which was plotted as a graph.

***Scratch Wound Assay***

MDA-MB-231 cells (40,000) were plated onto IncuCyte ImageLock 96-well cell culture microplates. After 18 hours, the cell monolayer was scraped using a wound-maker mechanical device (Essen BioScience), then washed with PBS and examined under inverted microscope. Scratch area was monitored using IncuCyte live-cell imaging system. Scratch-wound results were compiled from 3 wells with one scratch in each well. At the 24h time point, closure of the control scratch wound was observed.

***Transwell assay***

Invasion assay was performed using the Costar Transwell Invasion chamber (USA). Transwell inserts were rehydrated with serum-free DMEM for 1 hour and 0.5 mL of 1X $10^4$ of MDA-MB-231 cells were added on the upper chamber. 500µL of DMEM with 10% FBS (as chemoattractant) was added to the lower wells of the 24-well plate. The medium was discarded after 24 h. Non-migratory cells were removed with cotton-tipped swabs, and the lower surface of the insert was stained with 0.5% crystal fast violet. The cells were counted and captured under a Nikon Eclipse 80i microscope at 10× magnification.

***Cytokine measurement***

Measurement of IL-6 levels were performed by sandwich enzyme-linked immunosorbent assay (ELISA) using Human IL-6 mini ABTS ELISA development kit (Peprotech, New Jersey, USA).

#### *Supplementary Figures*

***Figure S1:*** (**A**) GPX8 expression significantly correlates with mesenchymal markers. Patients gene expression data was generated by the TCGA project and analyzed using the cBioportal webtool. The correlation between GPX8 and all the genome was calculated using the Bioportal website. The genes are ranked based on their Spearman correlation and subjected to Gene-set enrichment analysis (GSEA). This analysis resulted in a significant correlation with the "Hallmark Epithelial-Mesenchymal Transition″ gene set. The P-value was computed by GSEA. (**B**) GPX8 expression increased in the highly metastatic melanoma cell line A375-MA2. mRNA from A375 and A375-MA2 was isolated. The relative level of GPX8, as well as other indicated EMT markers in the two cell lines, were determined by quantitative real-time PCR (qPCR). Each value represents the mean ± SD for n=3.

***Figure S2:*** The expression of ZEB1, DPYD, and VIM in patients does not associate with poor patient prognosis: Kaplan-Meier survival plots for patients with breast cancer segregated based on GPX8 "high" versus "low" expression. The columns represent data from all breast cancers for the various genes as indicated.

***Figure S3:*** GPX8 KO does not affect proliferation. (**A**) MDA-MB-231 WT and GPX8 KO cells were infected with the indicated hairpins, and the proliferation rate was measured using CellTiterGlo. (**B**) GPX8 KO inhibits the expression of mesenchymal markers. HMLE-Twist-ER cells were infected and treated as indicated. The expression levels of the indicated genes were measured using qPCR (mean ± SEM). (**C**) Representative cell migration images of each indicated sample as generated by Incucyte, green color represents the initial scratch wound at time 0 hour and after the 24 hours the cells were migrated to the scratch area of the wound (brown color) . Bar = 600 µm.

***Figure S4:*** GPX8 loss in MDA-MB-231 cells affects cancer stemness. (**A**) representative images of mammosphere formations. (**B**) GPX8 loss affects tumor formation in mice. Female NOD-SCID mice were injected with $1X10^6$ cells generated from the different samples. After six weeks, the tumors were weighed; red circles represent mice without detectable tumors. (**C**) Tumors formation in mice. Upper image, 3 tumor samples taken out from mice injected with GPX8-WT cells. Lower image: 3 representative pictures of tumors taken out from mice injected with GPX8-WT cells, and bottom picture of the three tumors taken out from mice injected with GPX8-KO-KO1 cells.

**Khatib_Fig.S1**

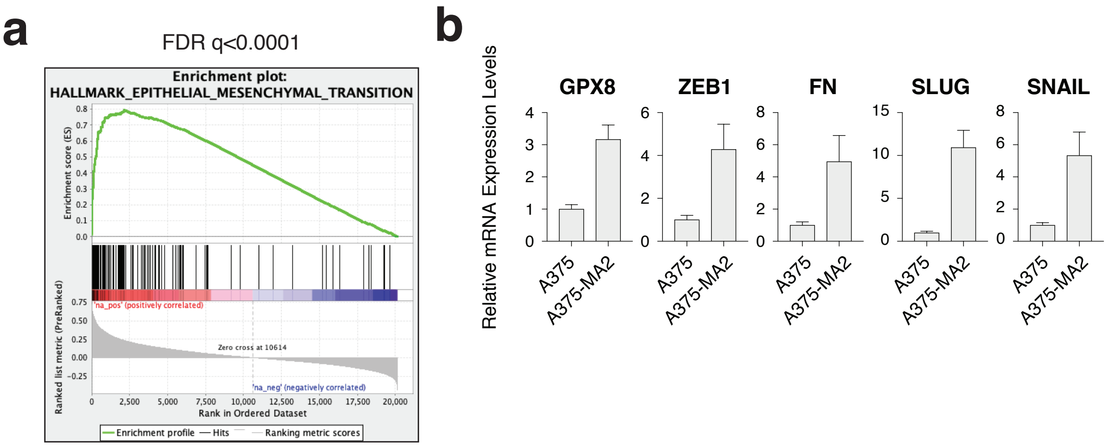

### Khatib_Fig.S2

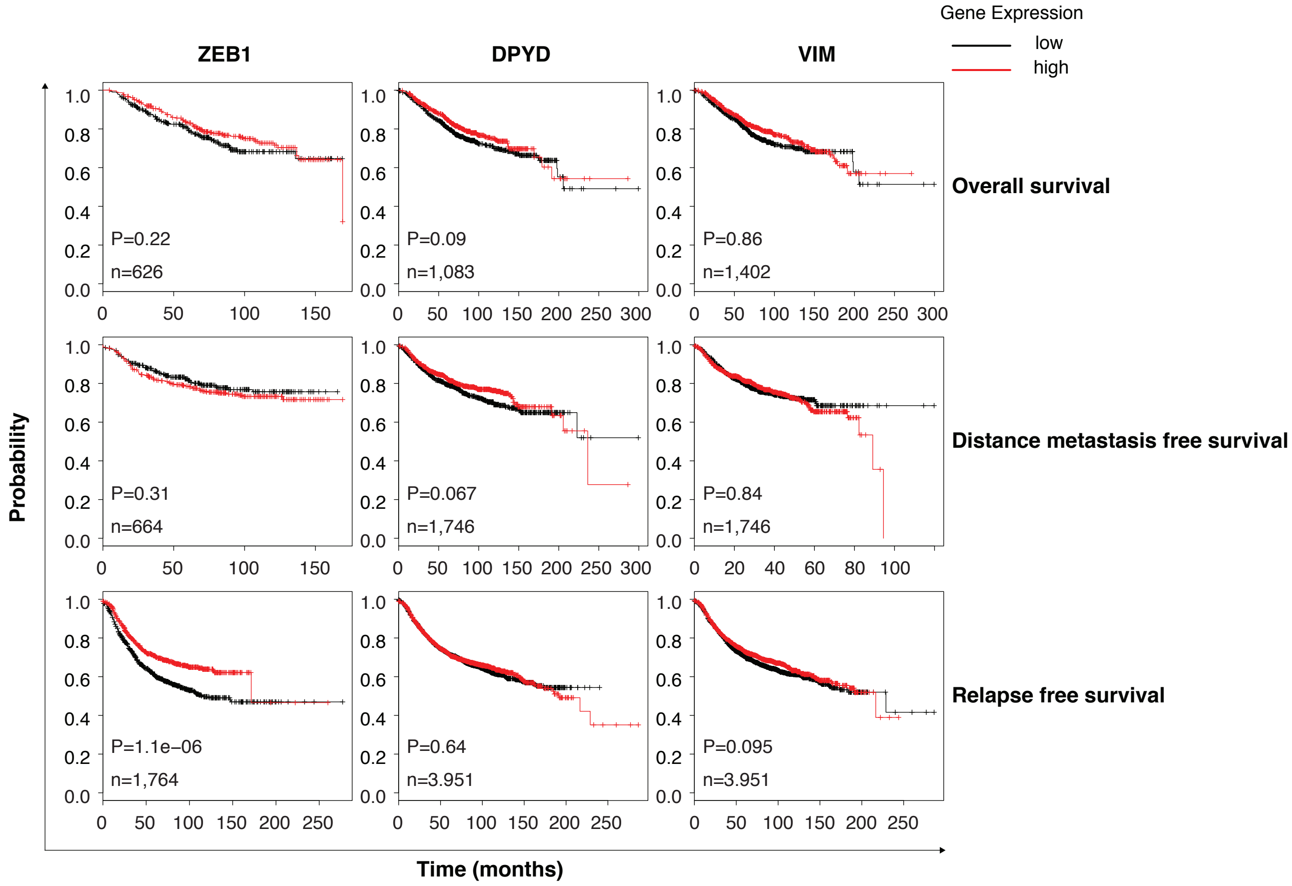

### Khatib_Fig.S3

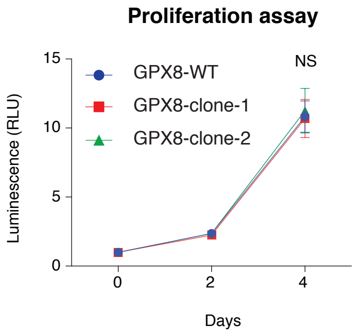

### Khatib_Fig.S4

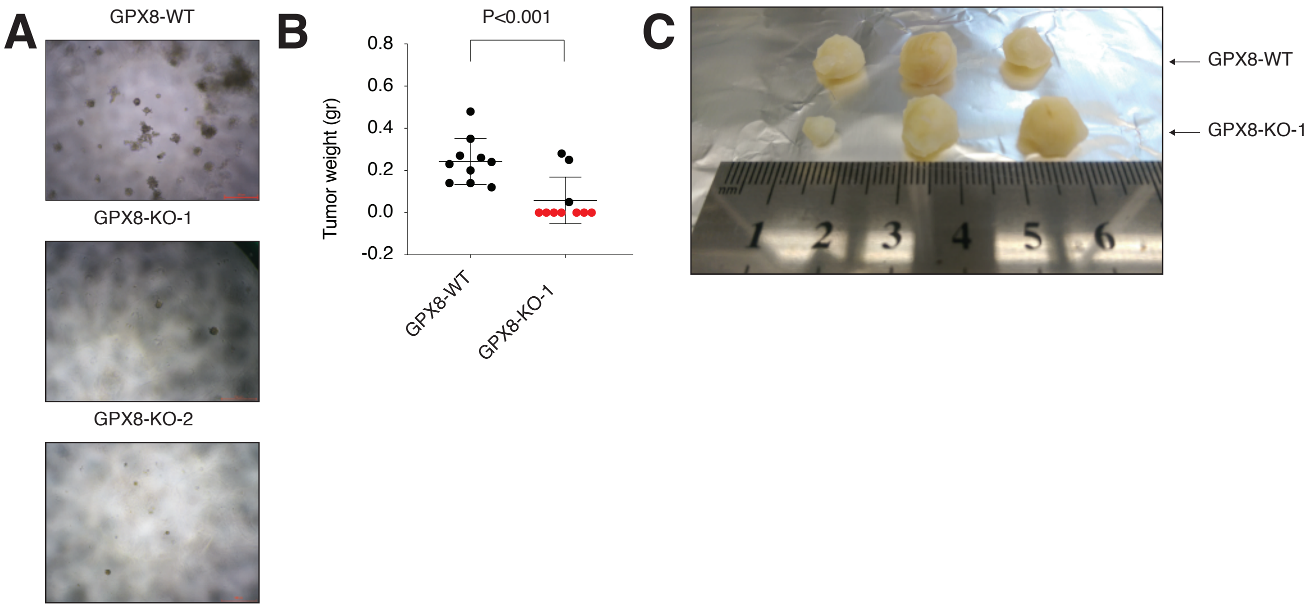

**Khatib_Fig.S5**

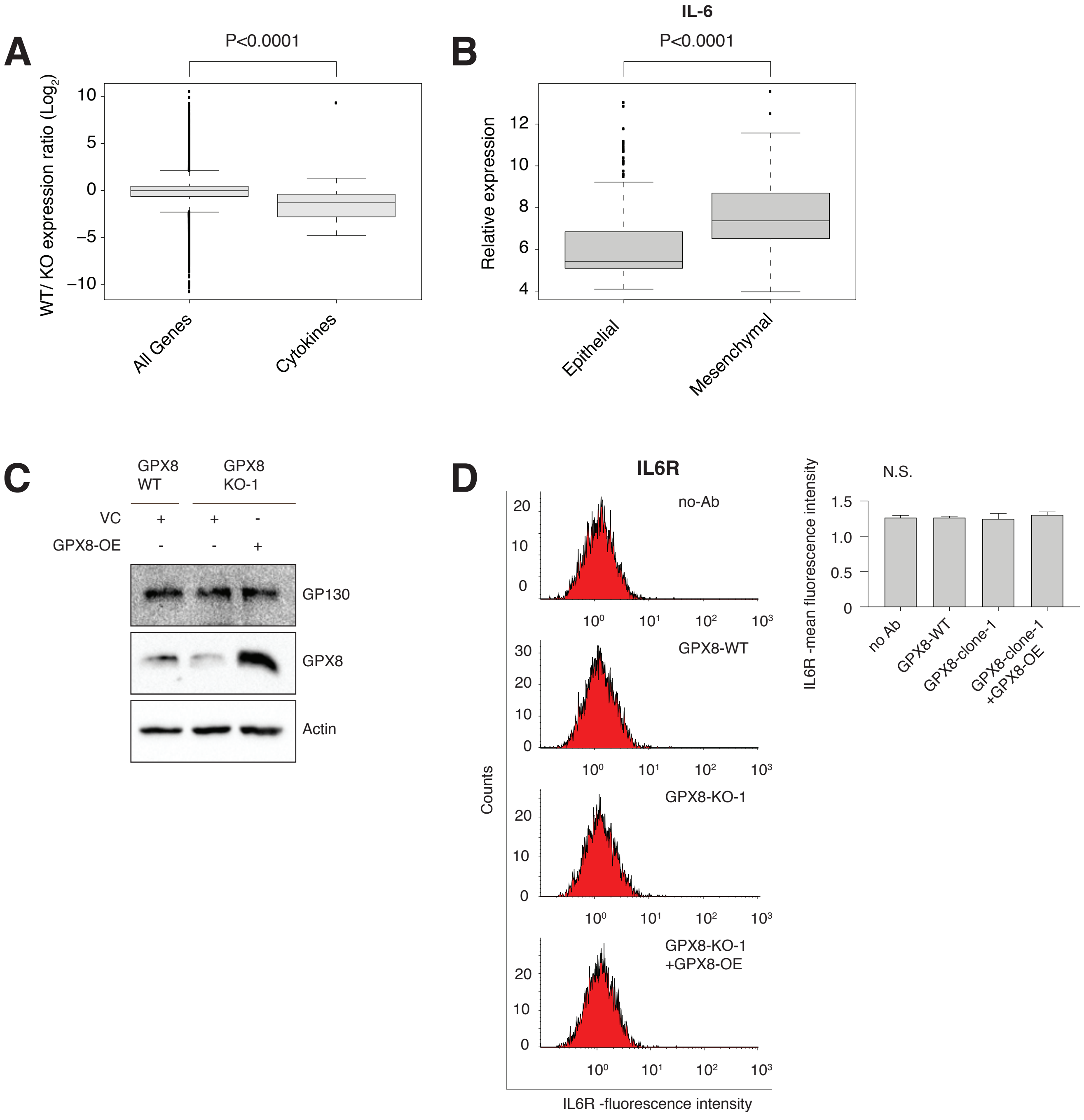

Table S1

| Cytokines | WT/GPX8 Expression ratio |
|---|---|
| TNFSF15 | -5.967651054 |
| CXCL11 | -4.795781986 |
| CXCL10 | -4.278606356 |
| IL1A | -3.311676461 |
| IL32 | -3.265509924 |
| CCL3 | -3.093808842 |
| CCL24 | -3.029077845 |
| CXCL16 | -2.945325114 |
| CSF3 | -2.91305564 |
| CXCL5 | -2.80683324 |
| IL1B | -2.665593976 |
| IL6 | -2.622305646 |
| IL15 | -2.586048629 |
| CCL5 | -2.530525858 |
| CSF1 | -2.423060392 |
| CSF2 | -2.312990936 |
| IL1RN | -2.244532193 |
| IL37 | -1.864472974 |
| TNFSF13 | -1.728147808 |
| LTA | -1.654497902 |
| IL34 | -1.632524105 |
| CCL20 | -1.577865526 |
| TGFB3 | -1.499327812 |
| CXCL1 | -1.311842852 |
| CCL28 | -1.205186878 |
| IFI27L2 | -1.177262465 |
| IL12B | -1.156854058 |
| LIF | -0.997461447 |
| TNFSF13B | -0.899610387 |
| CCL26 | -0.789560853 |
| TNF | -0.68451477 |
| IL4I1 | -0.660508031 |
| IL24 | -0.645971174 |
| CXCL3 | -0.617516266 |
| CXCL2 | -0.529600764 |
| CXCL8 | -0.405357891 |
| TNFSF12 | -0.371442046 |
| TGFB1 | -0.354676017 |
| IL18BP | -0.347809576 |
| IL11 | -0.049611741 |
| IL16 | 0.145442808 |
| IL23A | 0.163592069 |
| TGFB2 | 0.303254317 |
| IL7 | 0.448332371 |

Table S1

| | |
|---|---|
| EPO | 0.673763641 |
| IL18 | 0.743860811 |
| IL17D | 0.842027692 |
| XCL1 | 0.947804834 |
| CXCL14 | 1.302707956 |
| CCL2 | 9.307442163 |
| IL10 | Not Present in the seq results |
| IL13 | Not Present in the seq results |
| IL17A | Not Present in the seq results |
| IL17B | Not Present in the seq results |
| IL17C | Not Present in the seq results |
| IL17F | Not Present in the seq results |
| IL2 | Not Present in the seq results |
| IL20 | Not Present in the seq results |
| IL21 | Not Present in the seq results |
| IL22 | Not Present in the seq results |
| IL22RA2 | Not Present in the seq results |
| IL25 | Not Present in the seq results |
| IL26 | Not Present in the seq results |
| IL27 | Not Present in the seq results |
| IL3 | Not Present in the seq results |
| IL31 | Not Present in the seq results |
| IL4 | Not Present in the seq results |
| IL5 | Not Present in the seq results |
| IL9 | Not Present in the seq results |
| OSM | Not Present in the seq results |
| IFNA1 | Not Present in the seq results |
| IFNA10 | Not Present in the seq results |
| IFNA13 | Not Present in the seq results |
| IFNA14 | Not Present in the seq results |
| IFNA16 | Not Present in the seq results |
| IFNA17 | Not Present in the seq results |
| IFNA2 | Not Present in the seq results |
| IFNA21 | Not Present in the seq results |
| IFNA4 | Not Present in the seq results |
| IFNA5 | Not Present in the seq results |
| IFNA6 | Not Present in the seq results |
| IFNA7 | Not Present in the seq results |
| IFNA8 | Not Present in the seq results |
| IFNB1 | Not Present in the seq results |
| IFNE | Not Present in the seq results |
| IFNG | Not Present in the seq results |
| IFNL1 | Not Present in the seq results |
| IFNL2 | Not Present in the seq results |
| IFNL3 | Not Present in the seq results |
| IFNW1 | Not Present in the seq results |

Table S1

| | |
|---|---|
| TNFSF8 | Not Present in the seq results |
| CD70 | Not Present in the seq results |
| FASL | Not Present in the seq results |
| TNFSF18 | Not Present in the seq results |
| TNFSF10 | Not Present in the seq results |
| CXCL12 | Not Present in the seq results |
| CXCL13 | Not Present in the seq results |
| CXCL17 | Not Present in the seq results |
| CXCL6 | Not Present in the seq results |
| CXCL9 | Not Present in the seq results |
| CCL1 | Not Present in the seq results |
| CCL11 | Not Present in the seq results |
| CCL13 | Not Present in the seq results |
| CCL14 | Not Present in the seq results |
| CCL15 | Not Present in the seq results |
| CCL16 | Not Present in the seq results |
| CCL17 | Not Present in the seq results |
| CCL18 | Not Present in the seq results |
| CCL19 | Not Present in the seq results |
| CCL21 | Not Present in the seq results |
| CCL22 | Not Present in the seq results |
| CCL23 | Not Present in the seq results |
| CCL25 | Not Present in the seq results |
| CCL27 | Not Present in the seq results |
| CCL3L3 | Not Present in the seq results |
| CCL4 | Not Present in the seq results |
| CCL4L2 | Not Present in the seq results |
| CCL8 | Not Present in the seq results |
| XCL2 | Not Present in the seq results |
| CCL7 | Not Present in the seq results |
| CXCL4 | Not Present in the seq results |
| CXCL7 | Not Present in the seq results |

**Supplement Table-2. Primers Used for qRT-PCR, Related to Experimental Procedures**

| Gene | Forward seq | Reverse seq |
|---|---|---|
| GPX8 | ACTTCAGCGTGTTGGCTTTT | AGGCCTGATGACTTCAATGG |
| VIM | ACCCGCAC AACGAGAAGGT | ATTCTGCTGCTCCAGGAAGCG |
| CDH1 | TTGCACCGGTCGACAAAGGAC | TGGATTCCAGAAACGGAGGCC |
| CDH2 | TGTCGGTGACAAAGCCCCTG | AGGGCATTGGGATCGTCAGC |
| CLDN1 | CCTCCTGGGAGTGATAGCAAT | GGCAACTAAAATAGCCAGACCT |
| OCLN | ACAAGCGGTTTTATCCAGAGT | GTCATCCACAGGCGAAGTTAAT |
| β-Actin | CCAACCGCG AGAAGATGA | CCAGAGGCGTACAGGGATAG |
| ZEB1 | TGCACTGAGTGTGGAAAAGC | TGGTGATGCTGAAAGAGACG |
| ZEB2 | CAAGAGGCGCAAACAAGCC | GGTTGGCAATACCGTCATCC |
| CDH11 | AGAGAGCCCAGTACACGTTGA | TTGGCATGATAGGTCTCGTGC |
| FN1 | AGGAAGCCGAGGTTTTAACTG | AGGACGCTCATAAGTGTCACC |
| SNAI2 (Slug) | GAGAATGGACCTGCAAGCCCA | AGTGCAAGTGATGCGTCCGC |
| SNAI1 | CTGGGTGCCCTCAAGATGCA | CCGGACATGGCCTTGTAGCA |
| NGFR | CCTACGGCTACTACCAGGATG | CACACGGTGTTCTGCTTGT |
| CSF3 | GCTGCTTGAGCCAACTCCATA | GAACGCGGTACGACACCTC |
| IL6 | ACTCACCTCTTCAGAACGAATTG | CCATCTTTGGAAGGTTCAGGTTG |
| IL15 | TTGGGAACCATAGATTTGTGCAG | GGGTGAACATCACTTTCCGTAT |
| CXL10 | GTGGCATTCAAGGAGTACCTC | TGATGGCCTTCGATTCTGGATT |
| IL1B | ATGATGGCTTATTACAGTGGCAA | GTC GGAGATTCGTAGCTGGA |

*All primers are against the human genes
